# Comprehensive characterization of the mutational landscape in localized anal squamous cell carcinoma

**DOI:** 10.1101/796367

**Authors:** Lucía Trilla-Fuertes, Ismael Ghanem, Joan Maurel, Laura G-Pastrián, Marta Mendiola, Cristina Peña, Rocío López-Vacas, Guillermo Prado-Vázquez, Elena López-Camacho, Andrea Zapater-Moros, Victoria Heredia, Miriam Cuatrecasas, Pilar García-Alfonso, Jaume Capdevila, Carles Conill, Rocío García-Carbonero, Karen E. Heath, Ricardo Ramos-Ruiz, Carlos Llorens, Ángel Campos-Barros, Angelo Gámez-Pozo, Jaime Feliu, Juan Ángel Fresno Vara

**Affiliations:** Biomedica Molecular Medicine SL, C/ Faraday 7, 28049, Madrid, Spain; Medical Oncology Department, Hospital Universitario La Paz, Paseo de la Castellana 261, 28046, Madrid, Spain; Medical Oncology Department, Hospital Clinic of Barcelona, Translational Genomics and Targeted Therapeutics in Solid Tumors Group, IDIBAPS, University of Barcelona, Carrer de Villarroel 170, 08036, Barcelona, Spain; Pathology Department, Hospital Universitario La Paz, Paseo de la Castellana 261, 28046, Madrid, Spain; Molecular Pathology and Therapeutic Targets Group, Hospital Universitario La Paz-IdiPAZ, Paseo de la Castellana 261, 28046, Madrid, Spain; Biomedical Research Networking Center on Oncology-CIBERONC, ISCIII, Av. Monforte de Lemos 5, 28029, Madrid, Spain; Molecular Oncology & Pathology Lab, Institute of Medical and Molecular Genetics-INGEMM, Hospital Universitario La Paz-IdiPAZ, Paseo de la Castellana 261, 28046, Madrid, Spain; Translational Oncology Lab, Hospital Universitario La Paz-IdiPAZ, Paseo de la Castellana 261, 28046, Madrid, Spain; Pathology Department, Hospital Clínic Universitari de Barcelona, Carrer de Villarroel 170, 08036, Barcelona, Spain; Medical Oncology Department, Hospital General Universitario Gregorio Marañón, C/ Dr Esquerdo 46, 28007, Madrid, Spain; Medical Oncology Service, Vall Hebron University Hospital. Vall Hebron Institute of Oncology (VHIO), Passeig de la Vall d’Hebron 119-129, 08035, Barcelona, Spain; Radiotherapy Oncology Department, Hospital Clínic Universitari de Barcelona, Carrer de Villarroel 170, 08036, Barcelona, Spain; Medical Oncology Department, Hospital Universitario 12 de Ocubre, Instituto de Investigación Sanitaria Hospital 12 de Octubre (imas12), UCM, CNIO, CIBERONC, Av. Córdoba s/n, 28041, Madrid, Spain; Institute of Medical and Molecular Genetics, IdiPAZ, Hospital Universitario La Paz /& CIBERER, Unit 753; ISCIII, Paseo de la Castellana 261, 28046, Madrid, Spain; Genomics Unit Cantoblanco, Parque Científico de Madrid, C/ Faraday 7, 28049, Madrid, Spain; Biotechvana SL, Parque Científico de Madrid, C/ Faraday 7, 28049, Madrid, Spain; Cátedra UAM-Amgen, Universidad Autónoma de Madrid, Ciudad Universitaria de Cantoblanco, 28049, Madrid, Spain

**Keywords:** anal squamous cell carcinoma, exome sequencing, prognosis, personalized medicine

## Abstract

**Purpose:** Anal squamous cell carcinoma (ASCC) is a rare neoplasm. Chemo-radiotherapy is the standard of care, with no therapeutic advances achieved over the past three decades. Thus, a deeper molecular characterization of this disease is still necessary.

**Methods:** We analyzed 46 paraffin-embedded tumor samples from patients diagnosed with primary ASCC by exome sequencing. A bioinformatics approach focused in the identification of high impact genetic variants, which may act as drivers of oncogenesis, was performed. The relation between genetics variant and prognosis was also studied.

**Results:** The list of high impact genetic variants was unique for each patient. However, the pathways in which these genes are involved are well-known hallmarks of cancer, such as angiogenesis or immune pathways. Additionally, we determined that genetic variants in *BRCA2, ZNF750, FAM208B, ZNF599* and *ZC3H13* genes are related with poor disease-free survival in ASCC.

**Conclusion:** High impact genetic variants in ASCC affect pathways involved in cancer development, and may play a role in the etiology of the disease. Variants in *BRCA2, ZNF750, FAM208B, ZNF599* and *ZC3H13* genes seem to be related with prognosis in ASCC. Sequencing of ASCC clinical samples appears an encouraging tool for the molecular portrait of this disease.

## Introduction

Anal squamous cell carcinoma (ASCC) is a rare tumor. In 2019, an estimated 8,300 new cases will occur in the United States, representing approximately 2.5% of all gastrointestinal cancers ^1^.

Since the 1970s, the standard treatment has consisted of a combination of 5-fluorouracil (5FU) with mitomycin C or cisplatin and radiotherapy ^2,3^. Despite this treatment being very effective for early-stage tumors, the disease-free survival (DFS) rate in T3-T4 or N+ tumors ranges between 40-70% ^4,5^. Patients diagnosed with ASCC do not benefit from targeted therapy or immunotherapy. In addition, there is insufficient information on molecular prognostic or response prediction factors.

With the improvements in high-throughput molecular techniques, it is possible to study several variables instead of the classical gene-centered view. These technical advances allow for the study of multiple genetic alterations from clinical samples. Exome sequencing (ES) has contributed to the identification of new disease-causing genes and is now being incorporated into clinical practice ^6^. Since the first work reporting ES ^7^, numerous medical sequencing projects have faced the challenge of identifying molecular alterations related to rare diseases or cancers ^8^. The Cancer Genome Atlas (TCGA) is making huge strides in characterizing several tumor types by comprehensive molecular techniques. However, ASCC is not included because this project is focused on more frequent tumors.

Previous studies have analyzed metastatic or primary ASCC tumors by ES or even by gene panels in an attempt to describe the most frequent alterations in this disease. These studies established *PIK3CA* as a frequently mutated gene in ASCC ^9–12^. However, the exact relationship between genetic variations, phenotype and tumor evolution is currently unknown..

In this study, we analyzed 46 ASCC formalin-fixed paraffin-embedded (FFPE) samples. On the one hand, we characterized the main genetic variants present in these tumors and the main biological processes in which these genes are involved whilst on the other hand, we identified those genes in which the presence of a genetic variant is associated with DFS in ASCC.

## Materials and methods

### Patients

Forty-six treatment-naive FFPE samples from patients diagnosed with localized ASCC were analyzed by ES. All tumor samples were reviewed by an experienced pathologist. All the samples contained at least 70% invasive tumor cells. Informed consent was obtained for all patients and the study was approved by the Hospital Universitario La Paz Research Ethics Committee. Patients were required to have a histologically-confirmed diagnosis of ASCC; be 18 years of age or older; have an Eastern Cooperative Oncology Group performance status score (ECOG-PS) from 0 to 2; have not received prior radiotherapy or chemotherapy for this malignancy, and present with no distant metastasis. Demographic characteristics related to the tumor and the treatments were collected. The presence of human papillomavirus (HPV) infection was determined using CLART^®^ HPV2 (Genomica).

### DNA isolation

One 10 mm section from each FFPE sample was deparaffinized and DNA was extracted with GeneRead DNA FFPE Kit (Qiagen), following the manufacturer’s instructions. Once eluted, DNA was frozen at −80 °C until use.

### Library preparation, exome capture and Illumina sequencing

ES from 46 FFPE samples of ASCC was performed. Purified DNA was quantified by Picogreen and mean size was determined by gel electrophoresis. Genomic DNA was fragmented by mechanical means (Bioruptor) to a mean size of approximately 200 bp. Then, DNA samples were repaired, phosphorylated, A-tailed and ligated to specific adaptors, followed by PCR-mediated labeling with Illumina-specific sequences and sample-specific barcodes (Kapa DNA library generation kit).

Exome capture was performed using the VCRome system (capture size of 37 Mb, Roche Nimblegen) under a multiplexing of 8 samples per capture reaction. Capture was strictly carried out following manufacturer’s instructions. After capture, libraries were purified, quantified and titrated by Real Time PCR before sequencing. Samples were then sequenced to an approximate coverage of 4.5 Gb per sample in Illumina-NextSeq NS500 (Illumina Inc.) using 150 cycles (2×75) High Output cartridges.

### Bioinformatics exome sequencing data processing

The quality of the ES experiments was verified using FASTQC (http://www.bioinformmatics.babraham.ac.uk/projects/fastqc).

First, adaptors were removed using Cutadapt ^13^, and FASTQ files were filtered by quality using PrinSeq ^14^; both tools are included in the GPRO Suite (Biotechvana) ^15^. Alignment of the sequences was achieved using the human genome h19 as the reference genome. The tools BWA ^16^, Samtools ^17^ and Picard Tools (http://picard.sourceforge.net) were used. Variant calling was performed using the MuTect tool from the GATK4 package ^18^ combined with PicardTools, first, to create a panel of normal samples (PON) and, second, for the variant calling ^19^. The PON was built using 11 samples from Iberian exomes from 1000 genomes (http://www.ncbi.nlm.nih.gov/sra/) and it was used to discard germline variants.

### Prioritization of high impact genetic variants

With the aim of establishing the genetic variants that may act as drivers of the disease, the VarSEQ™ software (Golden Helix) was used. Variant Call Format (VCF) files were filtered according to strict criteria: a reading depth of at least 15x, a gnomAD global frequency <1%, high impact (frameshift, splice variants, stop-loss and stop-gain variants), and a detectable presence in at least 15% of the reads ^20^.

### Filtering of the most frequent genetic variants in our cohort

On the other hand, with the aim of establish the most frequent high and moderate impact variants, genetic variants were further annotated using Variant Effect Predictor (VEP) ^21^ and the Varsome database (https://varsome.com/). Then, the information provided by VEP was used to filter the genetic variants. The filtering criteria were: a frequency in the general population, according to the gnomAD database, of less than 1%, a high or moderate impact, and the presence of the variant in at least 10% of the patients in our cohort.

### Visual validation of the high impact genetic variants

The BAM files containing the prioritized variants were verified using Alamut Visual v2.11 (Interactive Biosoftware).

### Pathway annotation

Pathway annotation of the genes containing the prioritized variants was done using PANTHER database (http://www.pantherdb.org/) and DAVID webtool ^22^ selecting GO-BP as the category and *Homo sapiens* as the background.

### Genes associated with prognosis

From the list of genes with high or moderate impact variants identified by VEP, those genes in which the presence of a genetic variant was associated with DFS were identified using BRB Array Tools, developed by Dr. Richard Simon’s team ^23^. Genes associated with DFS were selected according to their p-values by a Kaplan-Meier analysis. DFS was defined as the time from primary tumor surgery until local and/or distant tumor relapse. For survival analyses, only the 41 patients treated with chemo-radiotherapy were included (Figure 1).

**Figure 1:**
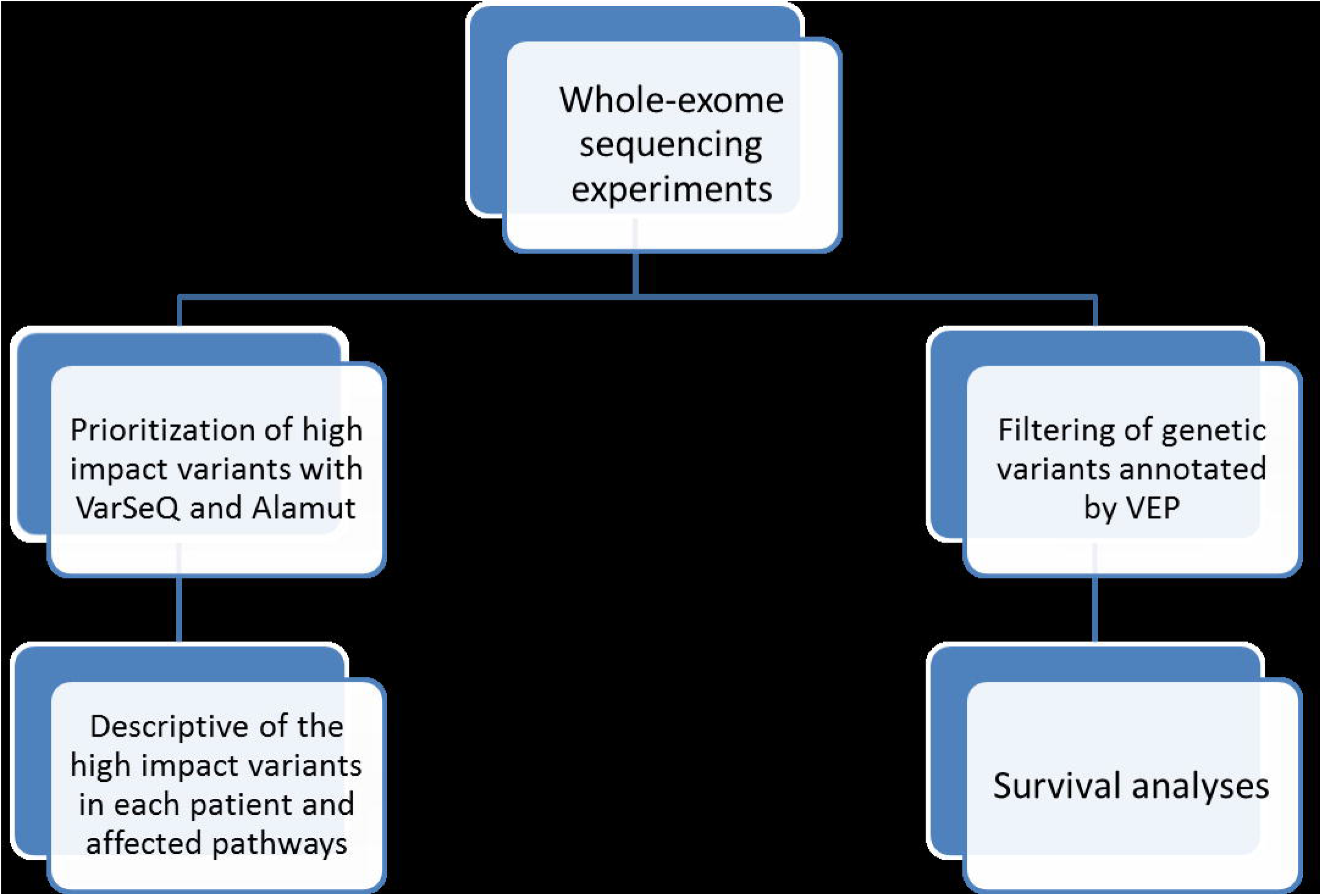
Workflow followed in this study.

### Statistical analyses

Statistical analyses were performed using GraphPad Prism v6 and IBM SPSS Statistics v20. All p-values were two-tailed and statistically significant was set as 0.05 or less.

## Results

### Patient cohort

Forty-six FFPE samples of ASCC were analyzed by ES. Twenty-eight samples were from patients included in the VITAL clinical trial (GEMCAD-09-02, NCT01285778). These patients were treated with panitumumab, 5FU and mitomycin C, concomitantly with radiotherapy. The remaining 18 patients were retrospectively included from the clinical practice in Hospital Universitario La Paz and Hospital Clinic. Fourteen patients were treated with cisplatin-5FU or mitomycin C-5FU, concomitantly with radiotherapy. Three patients, who were initially treated only with surgery and one patient treated with radiotherapy alone, were excluded from the survival analyses (Table 1).

**Table 1:**
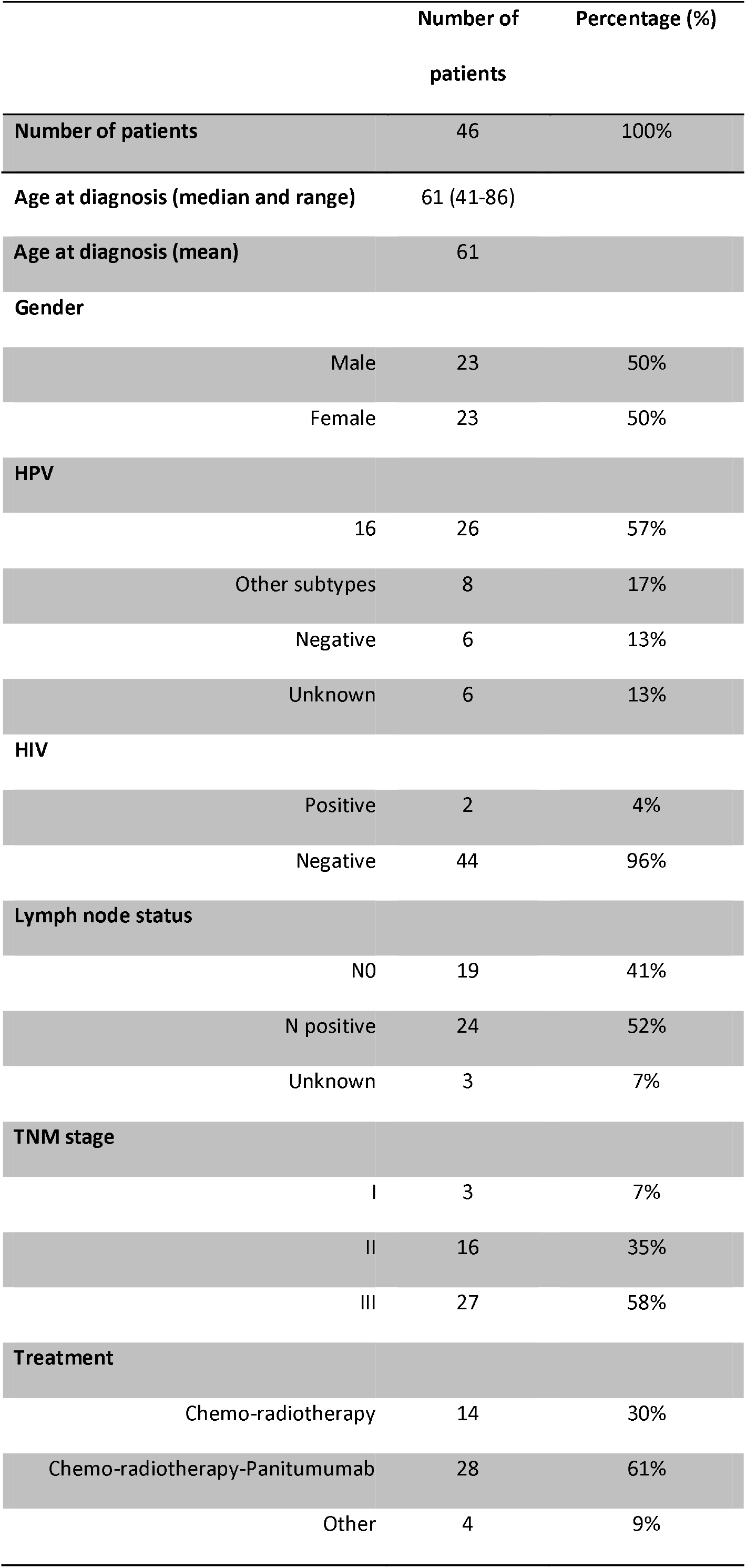
Patient clinical characteristics.

### Exome Sequencing experiments

The mean coverage obtained in ES experiments was > 42.6x, with the exception of one sample, with a coverage of 3.57x that was excluded from subsequent analyses. Once this sample was dismissed, the remaining samples presented a mapping efficiency of between 90-98% with the exception of one sample (75.4%). The human exome has > 195,000 exonic regions, out of which only 23,021 (11.21%) were not mapped in any sample.

### Relevant genes and their associated genetic variants

VarSEQ and Alamut were used to filter and visualize the variants that may play a significant role in the development of ASCC. For this purpose, we studied all genetic variants that caused a high impact in their respective gene. A total of 333 high impact variants across 312 genes in the 45 patients were prioritized by the VarSEQ software.

The list of the implicated genes with high impact variants is shown in Supp. Table 1.

Within the high impact variants, the most frequent type of alteration was the nonsense substitution which introduces a premature STOP codon (Table 2).

**Table 2:** Description of the type of high impact alterations detected by ES analysis.

| Type of alteration | Frequency |
| --- | --- |
| Duplications | 16 (4.8%) |
| Insertions | 32 (9.6%) |
| Deletions | 55 (16.4%) |
| Nonsense substitutions | 204 (62%) |
| Splice site substitutions | 19 (6%) |
| Intronic deletions | 2 (0.6%) |
| Intronic substitutions | 2 (0.6%) |

The principal alterations identified by these analyses were different between patients; i.e. each patient presented a unique set of high impact genetic variants. However, in some cases, the genes affected by these high impact alterations were common. The list of these genes is shown in Table 3 and Figure 2.

**Table 3:**
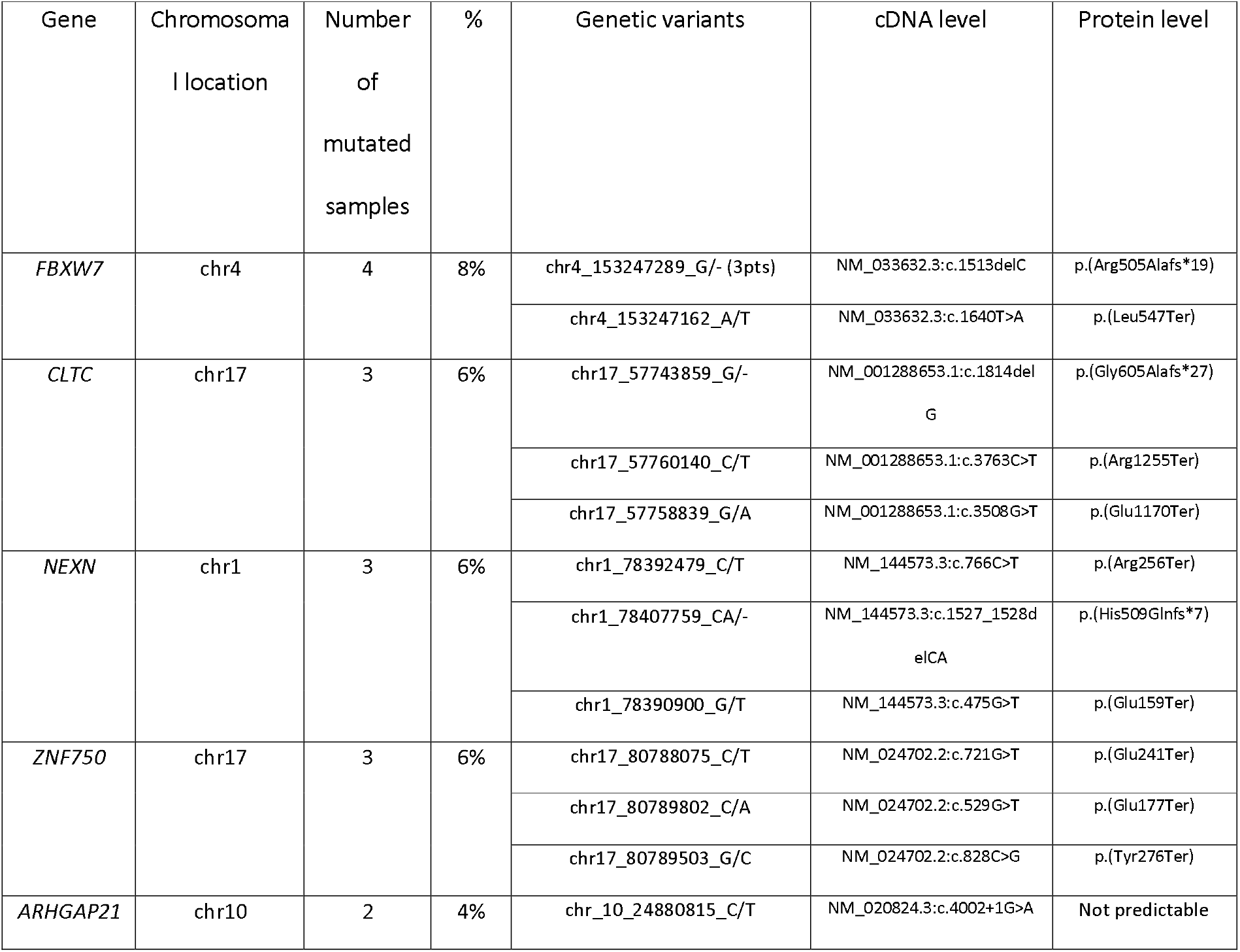
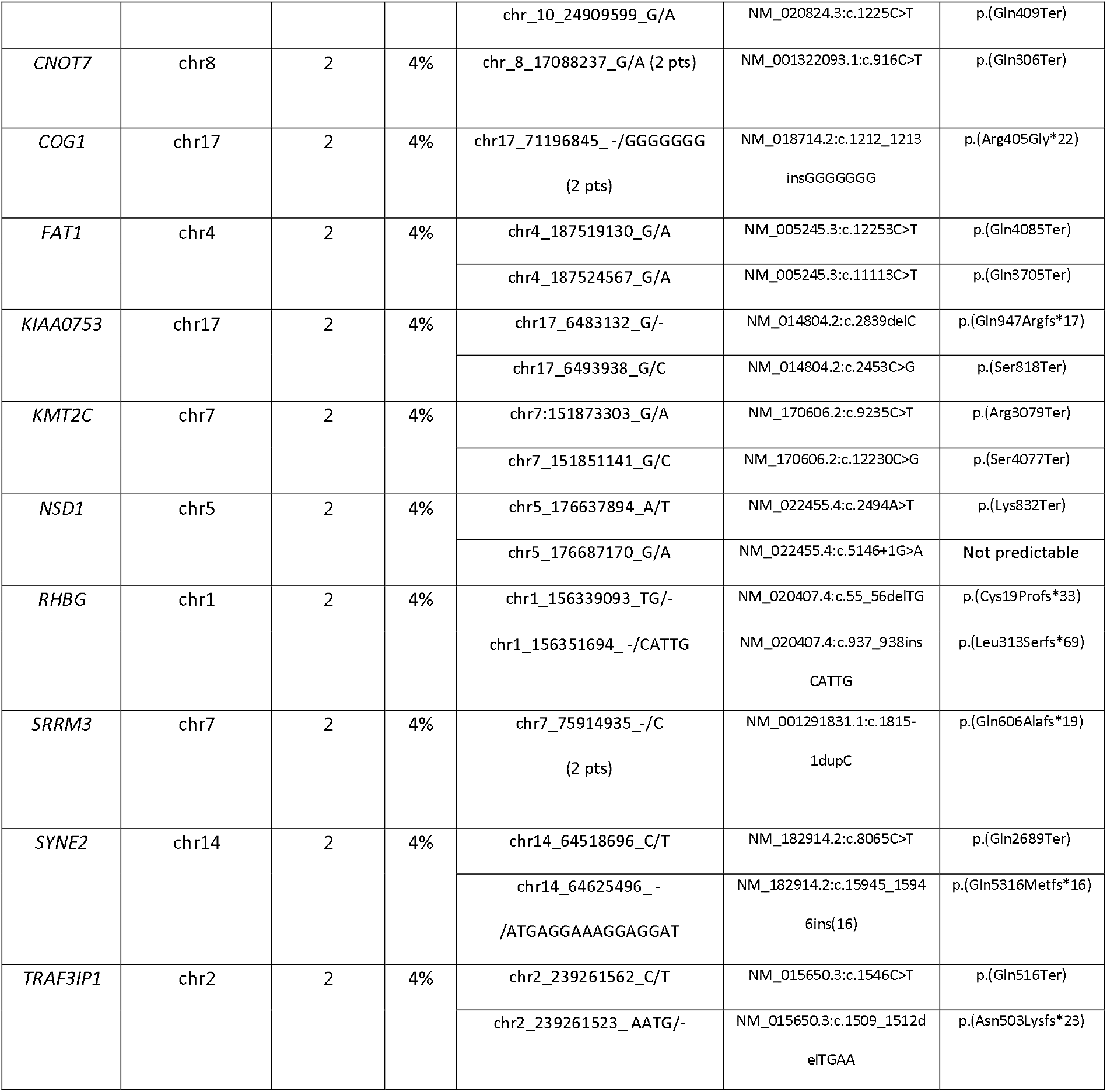
High impact genetic variants identified in more than one patient.

**Figure 2:**
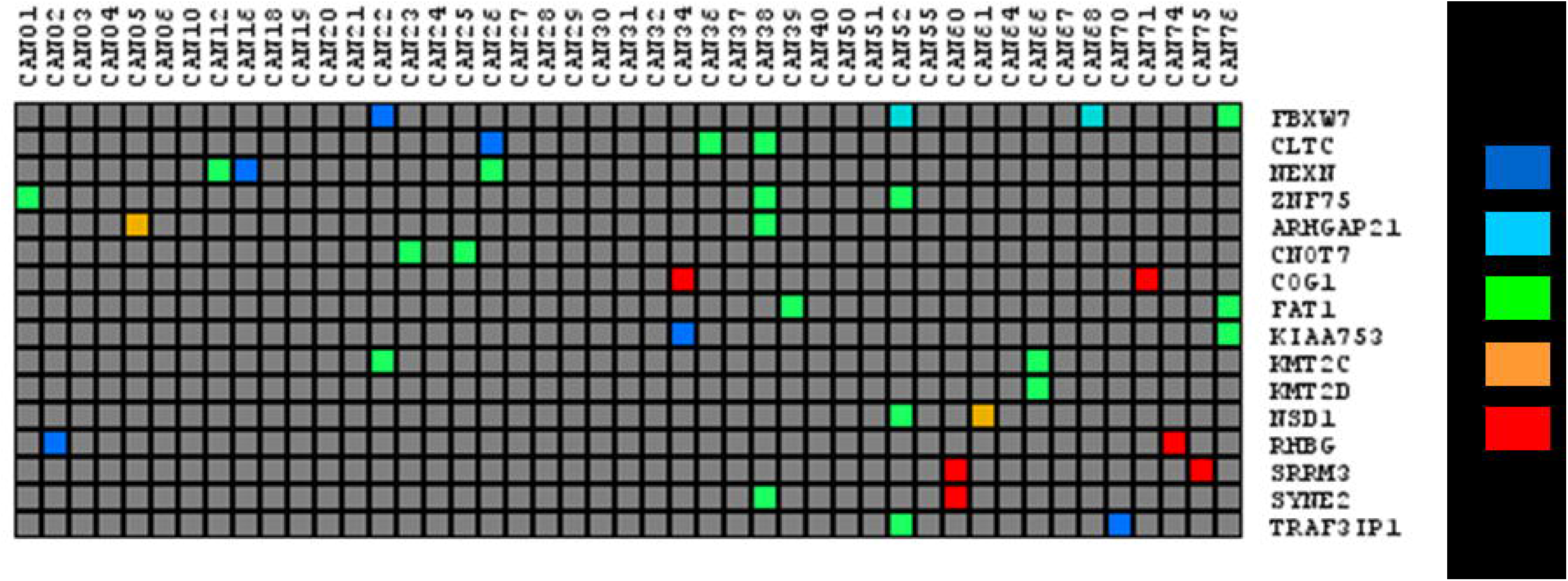
List of high impact genetic variants identified by VarSEQ in our cohort in more than one patient.

### Ontology analysis of the high impact genetic variants identified

With the aim of identifying the pathways affected by the genetic alterations, PANTHER and DAVID databases were used. The most frequently implicated pathways were intracellular signal transduction (22% of the genes, 24% of the patients in the cohort), immune (20% of the genes, 15% of the patients), and apoptosis (17% of the genes, 15% of the patients) pathways, although there were also relatively frequent alterations in genes implicated in angiogenesis (11% of the genes, 11% of the patients), metabolism (11% of the genes, 9% of the patients), chromatin modification (8% of the genes, 11% of the patients), EGFR signaling (8% of the genes, 7% of the patients) or Wnt signaling pathways (8% of the genes, 11% of the patients) (Figure 3).

**Figure 3:**
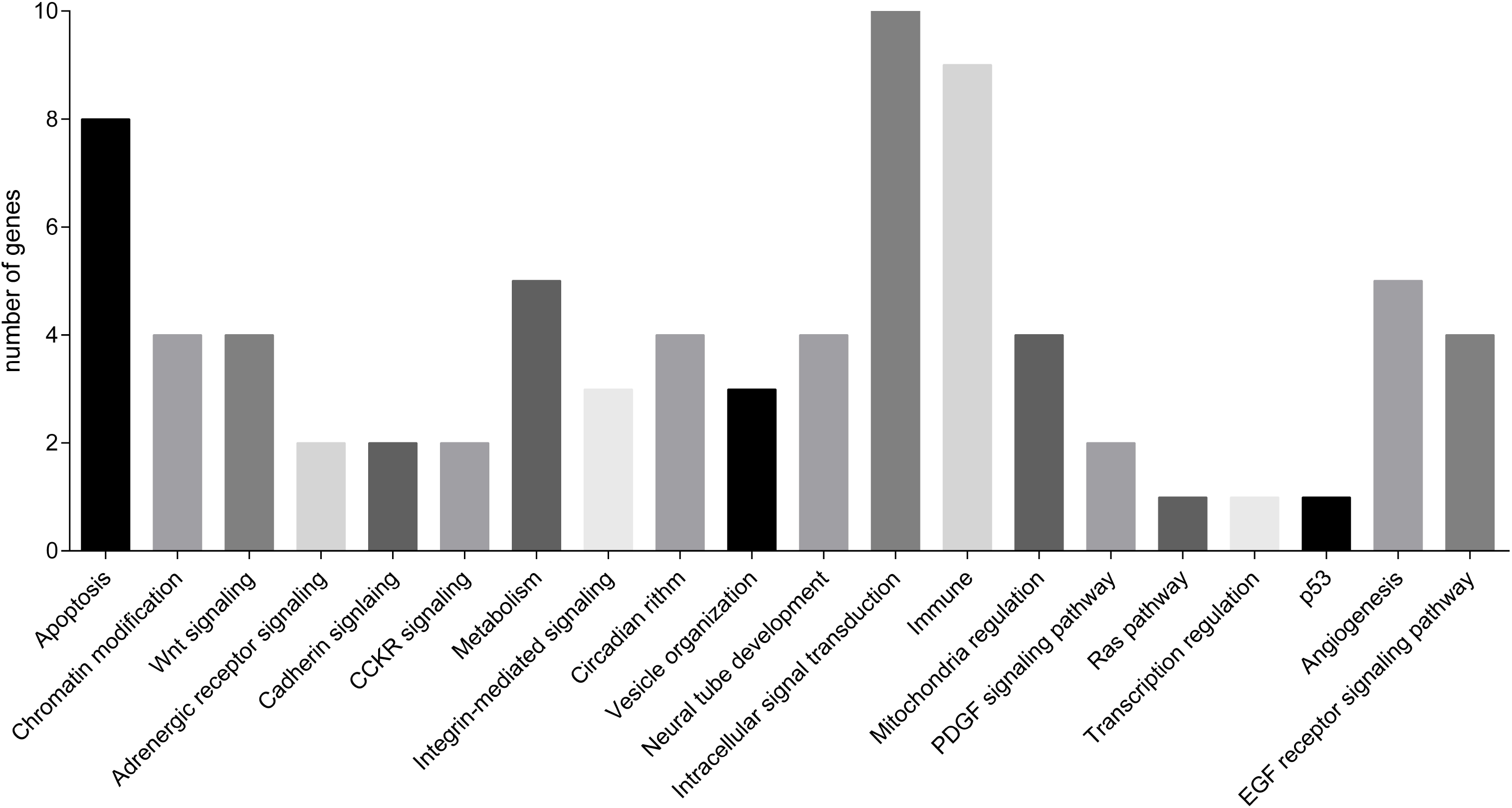
Pathways in which genes presenting high impact genetic variants are involved.

### Characterization of most frequently mutated genes in our cohort

With the aim of characterizing the most frequently mutated genes in ASCC, genetic variants were annotated using VEP and genes with high and moderate impact genetic variants were analyzed. After filtering by VEP results, 382 genes were found to present at least one high or moderate impact genetic variant in at least 10% of the patients in our cohort (Supp. Table 2). The functions associated with these 382 genes were cytoskeleton, DNA repair, adhesion and chromatin binding. *PIK3CA* was mutated in 40% of the patients, *FBXW7* in 16%, *FAT1* in 18%, and *ATM* in 27% of the patients. The most frequent variants found in *PIK3CA* are rs104886003 (7 patients), classified as a variant of uncertain significance (VUS) in Varsome, and rs121913273 (3 patients), classified as likely pathogenic by the same database.

### Genetic variants associated with prognosis in ASCC

With the aim of determining the genes associated with relapse in ASCC, a Kaplan-Meier analysis was performed. This analysis showed that in this cohort the presence of a high or moderate impact genetic variant in *BRCA2, ZNF750, FAM208B, ZNF599* and *ZC3H13* was associated with poor disease-free survival (Figure 4). The genetic variants detected in these genes are summarized in Table 4. Presenting more than one genetic variant in any of these genes implied a worse DFS (p=0.001) (Figure 5). Patients without genetic variants or with only a single genetic variant in these genes did not reach the median DFS and DFS percentages at 60 months were 54% and 29%, respectively. The median DFS in those patients with 2 or more genetic variant in these genes was 7 months and DFS percentage at 60 months was 86%.

**Figure 4:**
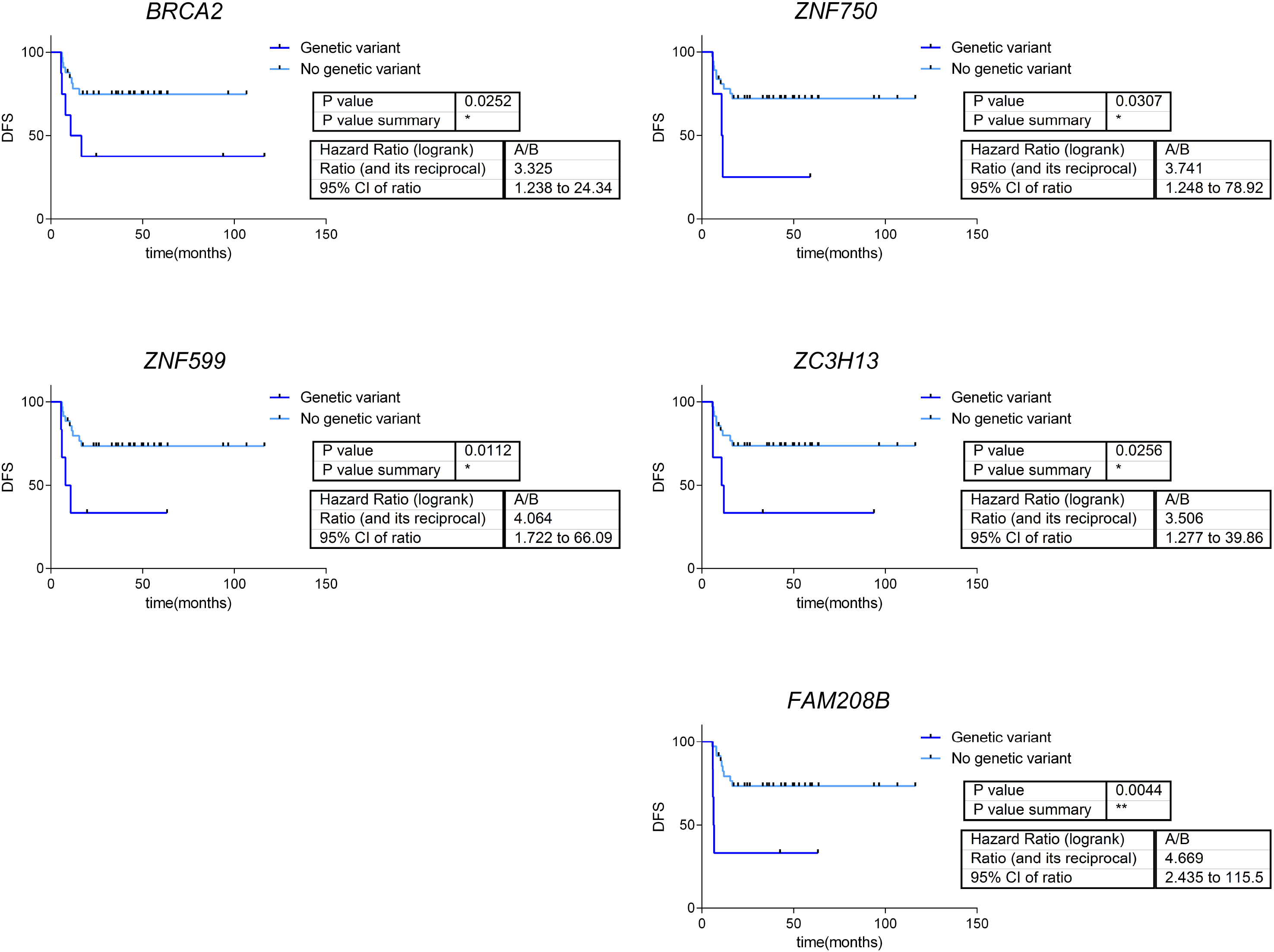
Survival curves for those patients whose high and moderate impact genetic variants are associated with disease-free survival (DFS).

**Table 4:**
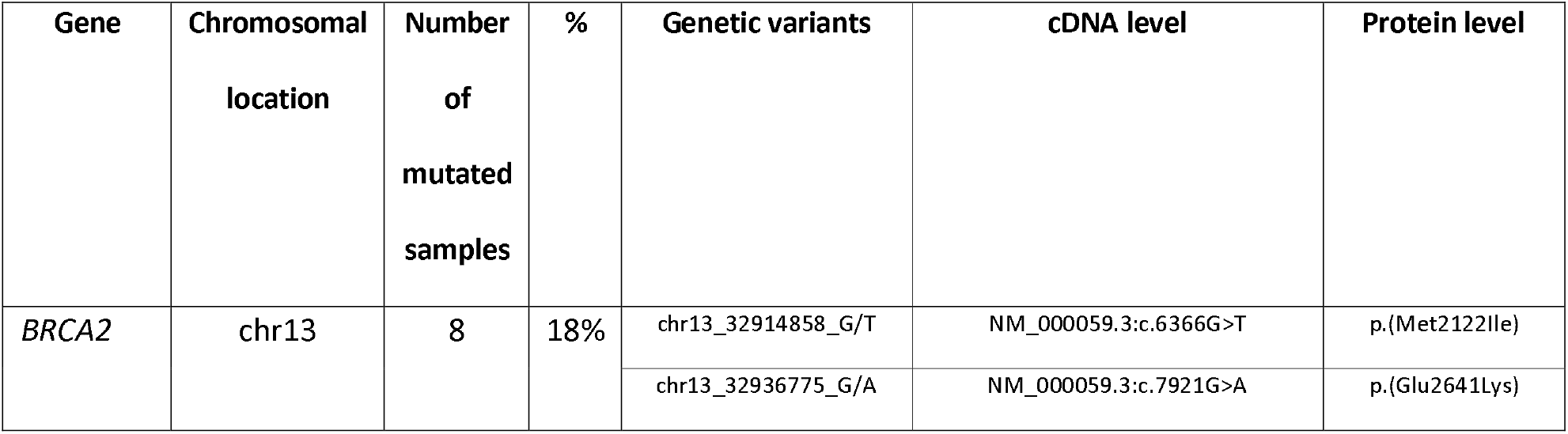
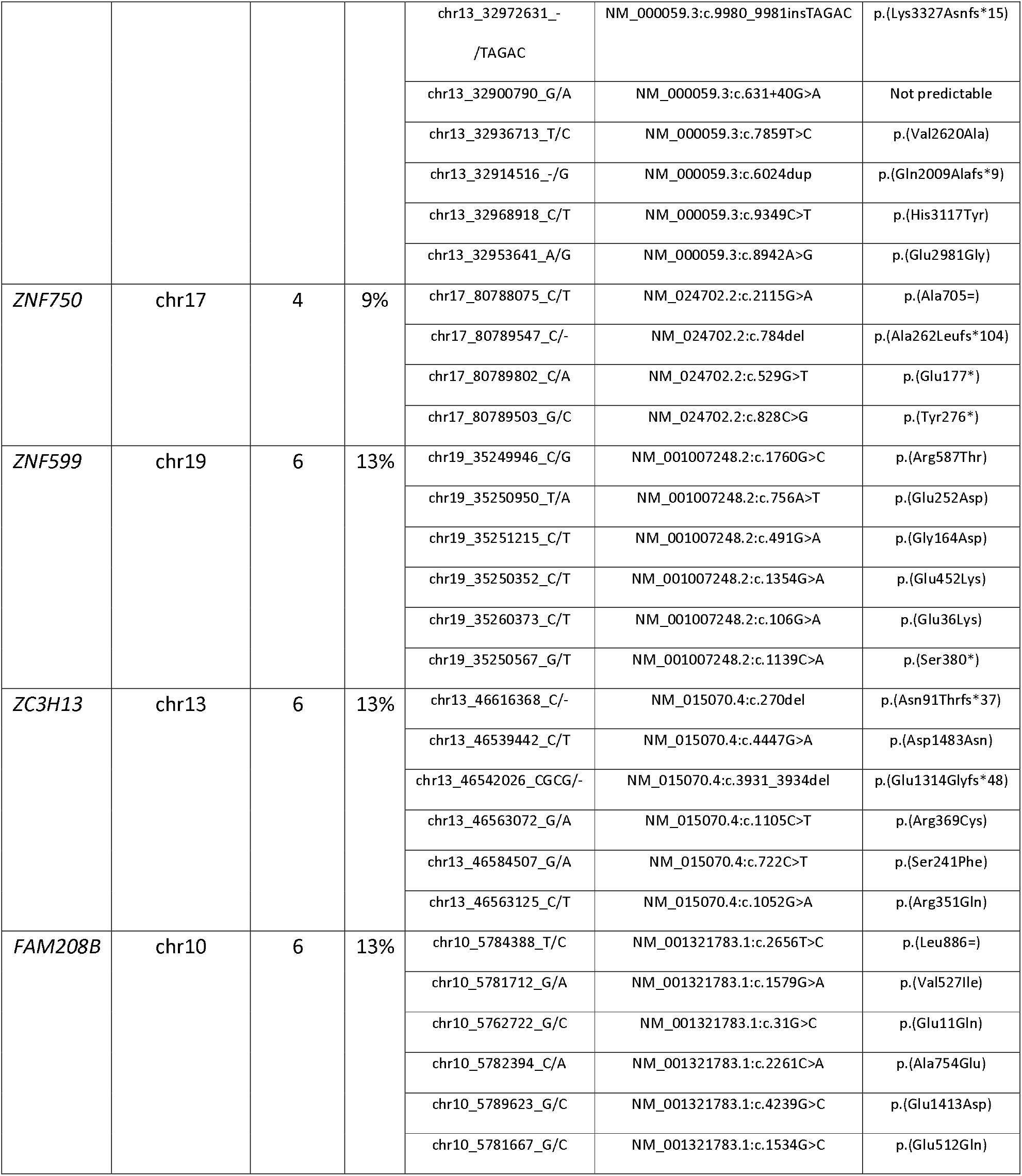
Genetic variants associated with disease-free survival in ASCC.

**Figure 5:**
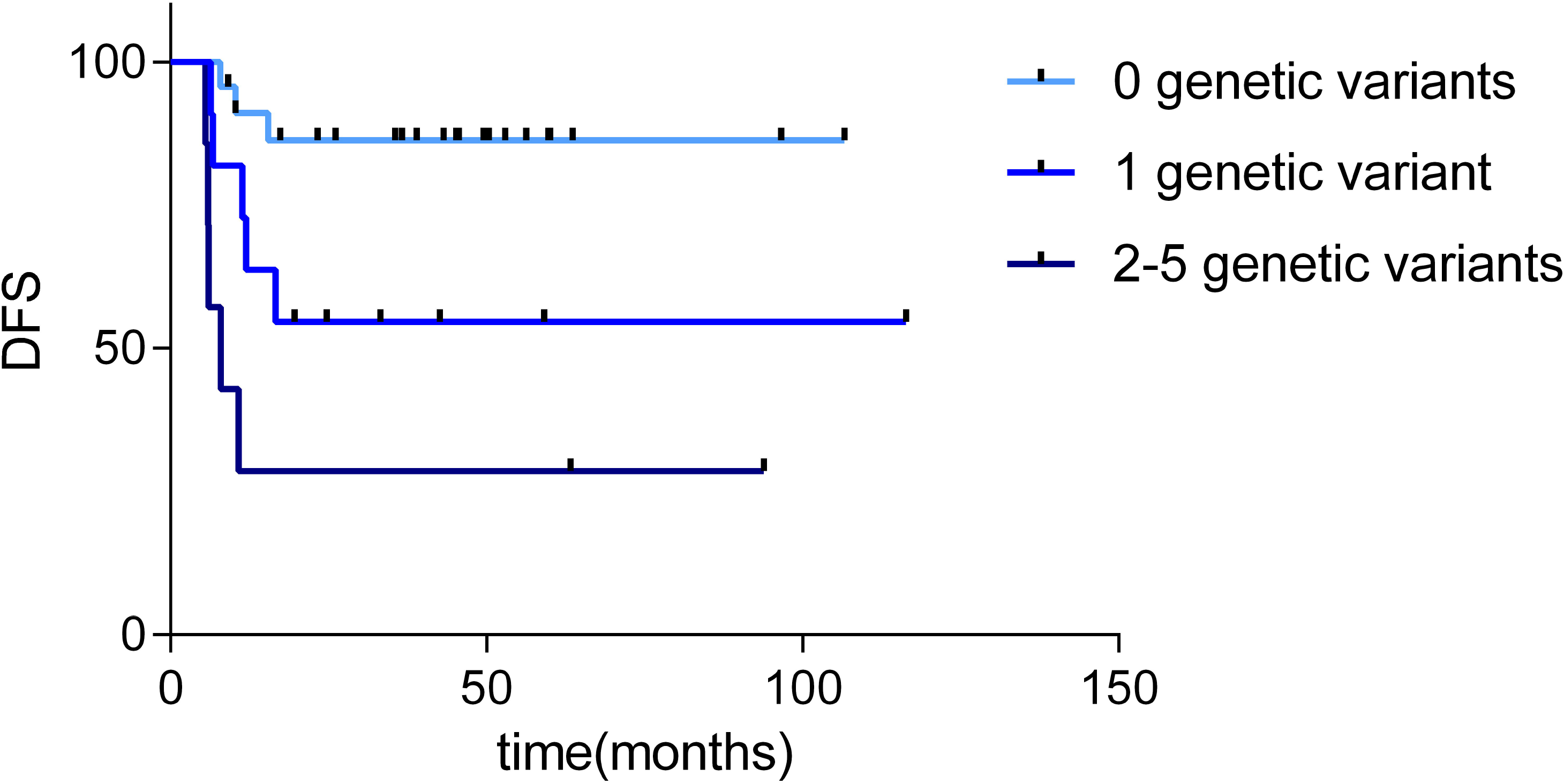
Survival curves obtained after considering the presence of high and moderate impact genetic variants in *BRCA2, FAM208B, ZNF750, ZNF599* and *ZC3H13* (0 genetic variants, n= 23 patients; 1 genetic variant, n= 11 patients; 2-5 genetic variants, n= 7 patients). DFS= disease-free survival.

## Discussion

ASCC is a relatively rare type of cancer, although its incidence has increased in recent years. The management of these patients consists of mitomycin C combined with 5FU chemotherapy and radiotherapy, but not all patients obtain a benefit from these treatments. For this reason, it is necessary to delve deeper into the molecular characterization of this disease, with the objective of identifying biomarkers and new potential therapeutic targets. In this study a cohort of 46 ASCC patients was studied by ES and a list of genes with high impact genetic variants that may be drivers of oncogenesis processes were identified. In addition, a characterization of the most frequently mutated genes with high and moderate impact variants was performed. Finally, five genes in which the presence of a genetic variant was associated with DFS were identified.

In this study, 46 ASCC paraffin samples were analyzed by ES. The analysis of FFPE samples using next-generation sequencing has been challenging due to the DNA fragmentation and the artificial alterations of the sequence caused by the fixation process. However, paired comparisons between fresh-frozen tissue and FFPE samples showed that although DNA damage in FFPE samples was evident, the results of fresh-frozen tissue and FFPE samples sequencing were comparable ^24^.

*PIK3CA* variants with a well-established functional effect were identified in 40% of our cohort samples. This value is consistent with that reported in previous studies ^11,12^. The most frequent *PIK3CA* variants were rs121913273 and rs104886003, both described in the COSMIC and Varsome databases. The rs121913273 variant has been described as likely pathogenic in esophageal and squamous cell carcinoma of the head and neck and in uterine cervical neoplasms (VCV000376244.2). Our results are comparable with the percentage observed in previous studies trying to identify the predominant genetic variants in ASCC ^9–12^. Cacheux et al. established variants in *PIK3CA* as a frequent alteration (20.3%) in this type of cancer ^10^. Chung et al. studied 70 patients and described recurrent alterations in 40 genes such as *PIK3CA* (40%), *FBXW7* (13%), *PTEN* (14%) and *RICTOR* (9%) ^12^. Both studies included localized, but also metastatic ASCC. Morris et al. analyzed metastatic ASCC samples combining ES and gene panels. They confirmed that *PIK3CA* is commonly mutated in ASCC (29%) and they used a xenograft model to test EGFR and PIK3CA inhibitors with a decrease in tumor growth ^11^. In a later work, Cacheux et al. performed ES on 20 ASCC patients and described *PIK3CA* (25%), *FBXW7* (15%), *FAT1* (15%) as frequently mutated genes, and *TRIP12* (15%), and chromatin remodeling as a pathway playing an important role in ASCC ^9^.

In contrast to the gene panel-based analyses, the non-directed ES approach made it possible to propose a list of genetic variants which may play an important role in the development of the disease, and to identify five genes in which the presence of a genetic variant is associated with DFS. Moreover, this analysis replicated the identification of genetic variants in some genes previously described in ASCC, such as *FBWX7,* but also suggested the involvement of novel genes, such as *ZNF750.*

In addition, we found that 16% of the patients presented a high or moderate impact variant in *FBXW7*, 18% in *FAT1*, and 27% in *ATM*. A remarkable finding was the presence of high impact genetic variants of the *FBXW7* gene in four patients. *FBXW7* is a cell cycle key regulator and its function as a tumor suppressor is well-known ^25^. Genetic variants in this gene have been previously described in 15% of ASCC patients. *FAT1*, which presented high impact variants in two patients of our cohort, has also been described as a tumor suppressor gene involved in the Wnt pathway ^26^. *FAT1* genetic variants have also been described in ASCC patients ^9^.

Strikingly, each patient presented a unique set of high impact variants. The most frequent alterations were nonsense stop-gain variants. However, the pathway analysis showed that, despite the individual variant diversity between patients, the affected biological pathways were shared between patients. This has already been seen in other tumors from The Cancer Genome Atlas studies ^27^.Moreover, the implicated pathways are traditionally associated with tumor progression and cancer development such as angiogenesis or metabolism pathways ^28^. The presence of high impact variants in the EGFR pathway in 7% of patients or in immunological processes may be relevant when proposing new therapies such as EGFR-targeted therapy or immunotherapy.

With the aim of determining the most frequent alterations in our cohort and identifying variants related to the progression of the disease, less restrictive filtering criteria were used. This led us to the identification of high or moderate impact genetic variants in 382 genes. These genes are mainly involved in DNA repair, chromatin binding, cytoskeleton and adhesion processes. Chromatin remodeling was previously suggested as a process with an important role in ASCC ^9^.

Survival analyses identified five genes associated with DFS. *ZNF750* was recently associated with prognosis in esophageal squamous cell carcinoma ^29,30^. Variants of *BRCA2* were also associated with esophageal squamous cell carcinoma and head and neck squamous cell carcinoma, both of which are related to HPV infection ^31,32^. Moreover, BRCA mutations are associated with response to PPAR inhibitors in pancreatic, breast and ovarian tumors ^33–35^. Therefore, the prevalence of *BRCA2* mutations may have therapeutic implications. Other genes associated with DFS (*FAM208B*, *ZNF599* and *ZC3H13*) had not been previously associated with cancer.

The two analyses performed offer complementary insights. The most restrictive filtering identified those genetic variants with a highly deleterious impact upon the resulting protein and proposed a list of variants that may be related to tumor development. In addition, based on the affected genes, the principal altered pathways involved were identified. In contrast, the alternative filtering pipeline yielded the most frequent variants in our cohort, some of them also associated with tumor progression and DFS.

ES analysis allowed us to identify candidate genes potentially involved in the etiology of ASCC without the bias of previous knowledge. On the other hand, our study led to the identification of genes that presented genetic variants associated with DFS in ASCC. As far as we know, this is the first time that an analysis of these characteristics has been performed. Another important point is the sample size. With ASCC being a rare tumor, it is difficult to gather a representative and homogeneous cohort. Our study analyzed 46 samples, the second largest cohort in ASCC from localized tumors treated with chemo-radiotherapy so far.

This study has some limitations. A validation with an independent cohort, especially for the prognostic genetic variants is still needed. Unlike the traditional genetic diseases, where an analysis pipeline is well-established, in tumor sequencing data, it is still necessary to establish a consensus analysis workflow.

To summarize, we analyzed 46 ASCC tumor samples by ES, an important cohort taking into account that ASCC is a rare tumor. Our study identified a set of variants having a high impact upon the corresponding protein and proposed a list of candidate genes which may play an important role in the etiology of ASCC. On the other hand, our study yielded a list of genetic variants apparently associated with poor disease progression. In conclusion, sequencing of ASCC clinical samples seems to be a promising tool for the molecular characterization of this pathology, which may aid the clinician in the prognostic stratification of patients and open new avenues for drug development.

## Acknowledgements

This study was supported by Instituto de Salud Carlos III, Spanish Economy and Competitiveness Ministry, Spain and co-funded by the FEDER program, “Una forma de hacer Europa” (PI15/01310), a Roche Farma funding, Amgen and a grant from Grupo Español Multidisciplinar en Cáncer Digestivo (GEMCAD1403). LT-F is supported by the Spanish Economy and Competitiveness Ministry (DI-15-07614). GP-V and EL-C are supported by the Consejería de Educación, Juventud y Deporte of Comunidad de Madrid (IND2017/BMD7783); AZ-M is supported by a Jesús Antolín Garciarena fellowship from IdiPAZ. The funders played no role in the study design, data collection and analysis, decision to publish or preparation of the manuscript.

## Supplementary files

Supp. Table 1: Genes with relevant high impact genetic variants according the prioritization criteria.

Supp. Table 2: 382 genes which presented a high or moderate impact genetic variant in at least 10% of the patients. 1= mutated; 0= non-mutated.

## References

1. Siegel RL, Miller KD, Jemal A. Cancer statistics, 2019. CA Cancer J Clin. 2019;69(1):7–34.

2. Benson A, Venook A, Al-Hawary M, et al. Anal carcinoma, Version 2.2018, NCCN Clinical Practice Guidelines in Oncology. In. Vol 7. J Natl Compr Canc Netw. 2018:852–871.

3. Glynne-Jones R, Nilsson PJ, Aschele C, et al. Anal cancer: ESMO-ESSO-ESTRO clinical practice guidelines for diagnosis, treatment and follow-up. Eur J Surg Oncol. 2014;40(10): 1165–1176.

4. Gunderson LL, Winter KA, Ajani JA, et al. Long-term update of US GI intergroup RTOG 98-11 phase III trial for anal carcinoma: survival, relapse, and colostomy failure with concurrent chemoradiation involving fluorouracil/mitomycin versus fluorouracil/cisplatin. J Clin Oncol. 2012;30(35):4344–4351.

5. Ajani JA, Winter KA, Gunderson LL, et al. Prognostic factors derived from a prospective database dictate clinical biology of anal cancer: the intergroup trial (RTOG 98-11). Cancer. 2010;116(17):4007–4013.

6. Tetreault M, Bareke E, Nadaf J, Alirezaie N, Majewski J. Whole-exome sequencing as a diagnostic tool: current challenges and future opportunities. Expert Rev Mol Diagn. 2015;15(6):749–760.

7. Ng S, Turner E, Robertson P, et al. Targeted capture and massively parallel sequencing of 12 human exomes. Nature. 2009;461 (7261):272–276.

8. Rabbani B, Tekin M, Mahdieh N. The promise of whole-exome sequencing in medical genetics. J Hum Genet. 2014;59(1):5–15.

9. Cacheux W, Dangles-Marie V, Rouleau E, et al. Exome sequencing reveals aberrant signalling pathways as hallmark of treatment-naive anal squamous cell carcinoma. Oncotarget. 2018;9(1):464–476.

10. Cacheux W, Rouleau E, Briaux A, et al. Mutational analysis of anal cancers demonstrates frequent PIK3CA mutations associated with poor outcome after salvage abdominoperineal resection. Br J Cancer. 2016;114(12):1387–1394.

11. Morris V, Rao X, Pickering C, et al. Comprehensive Genomic Profiling of Metastatic Squamous Cell Carcinoma of the Anal Canal. Mol Cancer Res. 2017;15(11):1542–1550.

12. Chung JH, Sanford E, Johnson A, et al. Comprehensive genomic profiling of anal squamous cell carcinoma reveals distinct genomically defined classes. Ann Oncol. 2016;27(7): 1336–1341.

13. Martin M. Cutadapt removes adapter sequences from high-throughput sequencing reads. EMBnetjournal. 2011;17(1): 10–12.

14. Schmieder R, Edwards R. Quality control and preprocessing of metagenomic datasets. Bioinformatics. 2011;27(6):863–864.

15. Futami R, Muñoz-Pomer A, Viu J, et al. GPRO The professional tool for annotation, management and functional analysis of omic databases. Biotechvana Bioinformatics: SOFT3. 2011.

16. Li H, Durbin R. Fast and accurate short read alignment with Burrows-Wheeler transform. Bioinformatics. 2009;25(14):1754–1760.

17. Li H, Handsaker B, Wysoker B, et al. The Sequence Alignment/Map format and SAMtools. Bioinformatics. 2009;25:2078–2079.

18. McKenna A, Hanna M, Banks E, et al. The Genome Analysis Toolkit: a MapReduce framework for analyzing next-generation DNA sequencing data. Genome Research. 2010;20(9): 1297–1303.

19. Cibulskis K, Lawrence M, Carter S, et al. Sensitive detection of somatic point mutations in impure and heterogeneous cancer samples. Nature Biotechnology. 2013;31(3):213–219.

20. Jalali Sefid Dasthi M, J G. A practical guide to filtering and prioritizing genetic variants. Biotechniques. 2017;62(1): 18–30.

21. Cunningham F, Amode MR, Barrell D, et al. Ensembl 2015. Nucleic Acids Res. 2015;43(Database issue):D662–669.

22. Huang dW, Sherman BT, Lempicki RA. Systematic and integrative analysis of large gene lists using DAVID bioinformatics resources. Nat Protoc. 2009;4(1):44–57.

23. Simon R. Roadmap for developing and validating therapeutically relevant genomic classifiers. J Clin Oncol. 2005;23(29):7332–7341.

24. Oh E, Choi YL, Kwon MJ, et al. Comparison of Accuracy of Whole-Exome Sequencing with Formalin-Fixed Paraffin-Embedded and Fresh Frozen Tissue Samples. PLoS One. 2015; 10(12):e0144162.

25. Takeishi S, Nakayama KI. Role of Fbxw7 in the maintenance of normal stem cells and cancer-initiating cells. Br J Cancer. 2014;111(6):1054–1059.

26. Morris LG, Kaufman AM, Gong Y, et al. Recurrent somatic mutation of FAT1 in multiple human cancers leads to aberrant Wnt activation. Nat Genet. 2013;45(3):253–261.

27. Bailey MH, Tokheim C, Porta-Pardo E, et al. Comprehensive Characterization of Cancer Driver Genes and Mutations. Cell. 2018;174(4):1034–1035.

28. Hanahan D, Weinberg RA. Hallmarks of cancer: the next generation. Cell. 2011; 144(5):646–674.

29. Otsuka R, Akutsu Y, Sakata H, et al. ZNF750 Expression Is a Potential Prognostic Biomarker in Esophageal Squamous Cell Carcinoma. Oncology. 2018;94(3):142–148.

30. Hazawa M, Lin DC, Handral H, et al. ZNF750 is a lineage-specific tumour suppressor in squamous cell carcinoma. Oncogene. 2017;36(16):2243–2254.

31. Akbari MR, Malekzadeh R, Lepage P, et al. Mutations in Fanconi anemia genes and the risk of esophageal cancer. Hum Genet. 2011;129(5):573–582.

32. Low GM, Thylur DS, Yamamoto V, Sinha UK. The effect of human papillomavirus on DNA repair in head and neck squamous cell carcinoma. Oral Oncol. 2016;61:27–30.

33. Munroe M, Kolesar J. Olaparib for the treatment of BRCA-mutated advanced ovarian cancer. Am J Health Syst Pharm. 2016;73(14):1037–1041.

34. Robson M, Goessl C, Domchek S. Olaparib for Metastatic Germline BRCA-Mutated Breast Cancer. N Engl J Med. 2017;377(18):1792–1793.

35. Golan T, Hammel P, Reni M, et al. Maintenance Olaparib for Germline BRCA-Mutated Metastatic Pancreatic Cancer. N Engl J Med. 2019.

