## Supplementary material for "Comprehensive characterization of the mutational landscape in localized anal squamous cell carcinoma": Sup Table 1

| *FBXW7* | *AQR* | *COQ5* | *FGFBP1* | *LILRA1* | *PA2G4* | *RPS6KA6* | *TDRD15* |
| --- | --- | --- | --- | --- | --- | --- | --- |
| *CLTC* | *ARHGAP45* | *CPN1* | *FITM1* | *LRP2* | *PCDHB14* | *RPS6KC1* | *TEP1* |
| *NEXN* | *ARID1A* | *CPNE3* | *FRRS1* | *MAP3K1* | *PDCD5* | *RTN3* | *TEX11* |
| *ZNF750* | *ARID1B* | *CREB3L1* | *FRY* | *MAP4K4* | *PDE4A* | *RYR2* | *TEX15* |
| *ARHGAP21* | *ARID5B* | *CREB5* | *FUNDC2* | *MAPK3* | *PDIA3* | *RYR3* | *TMEM38B* |
| *CNOT7* | *ARPC1A* | *CSDC2* | *GALNT5* | *MARCH6* | *PDZK1* | *SCAF4* | *TMIGD1* |
| *COG1* | *ASB10* | *CT83* | *GALNT7* | *MARCH7* | *PEAK1* | *SCFD2* | *TMX2* |
| *FAT1* | *ATP6V0E1* | *CTH* | *GCDH* | *MATR3* | *PGBD1* | *SCGN* | *TRHDE* |
| *KIAA0753* | *BAIAP2* | *CTNNA2* | *GCLC* | *MEDAG* | *PHF3* | *SCIN* | *TRIP12* |
| *KMT2C* | *BBOF1* | *CYP2R1* | *GIMAP4* | *MIS18BP1* | *PHTF1* | *SCN3A* | *TTC21A* |
| *KMT2D* | *BBS7* | *DCHS2* | *GLMN* | *MKNK1* | *PI4KB* | *SCNM1* | *TXLNG* |
| *NSD1* | *BCORL1* | *DDX58* | *GLS2* | *MLPH* | *PIK3R6* | *SCUBE2* | *TYROBP* |
| *RHBG* | *BOC* | *DEFB135* | *GNMT, PEX6* | *MPDU1* | *PKD2L1* | *SEPT10* | *U2AF2* |
| *SRRM3* | *BTNL3* | *DGKI* | *GNPAT* | *MPHOSPH8* | *PLCB4* | *SGO2* | *UBR4* |
| *SYNE2* | *C1orf210* | *DIAPH2* | *GPAT4* | *MPO* | *PLOD3* | *SH2D4B* | *UBR5* |
| *TRAF3IP1* | *C20orf194* | *DIEXF* | *GPRC5A* | *MPP1* | *PODN* | *SHANK1* | *UNC80* |
| *RBM43* | *C2CD3* | *DLG5* | *GPSM2* | *MRM3* | *PPWD1* | *SLC13A2* | *USH1C* |
| *A2M* | *C8B* | *DLX4* | *GPT* | *MRPL40* | *PREX2* | *SLC40A1* | *USP12* |
| *ABCA10* | *CACNA1G* | *DNAH10* | *GSDMA* | *MTERF3* | *PRKAG2* | *SLC9A7* | *VWA5B1* |
| *ABCC9* | *CCDC121* | *DNAH7* | *H2BFM* | *MYBPC1* | *PRKCD* | *SMURF1* | *VWDE* |
| *ACOXL* | *CCDC180* | *DNMT1* | *HELZ* | *MYH11* | *PRKCE* | *SNX18* | *WASF1* |
| *ACSM2A* | *CCDC82* | *DNPEP* | *HHAT* | *MYL6B* | *PRKD1* | *SNX29* | *WBSCR28* |
| *ACTR3* | *CCL5* | *DST* | *HIGD1B* | *MYO18B* | *PROK2* | *SNX32* | *YKT6* |
| *ADAM11* | *CCNA1* | *DZIP3* | *HIPK3* | *MYOF* | *PROX1* | *SORBS1* | *ZCWPW2* |
| *ADAM20* | *CD177* | *EGR4* | *HK3* | *NCOA2* | *PRR14* | *SORCS1* | *ZFP69B* |
| *ADCY4* | *CD5* | *EIF6* | *HLCS* | *NCOR1* | *PTEN* | *SPATA20* | *ZFYVE16* |
| *AGL* | *CDC37L1* | *ELF1* | *IGSF10* | *NDUFV1* | *PTPN14* | *SPATA31E1* | *ZFYVE26* |
| *AIM1* | *CDH7* | *EPHA2* | *IKBKB* | *NEB* | *PUM2* | *ST6GALNAC3* | *ZNF200* |
| *AK5* | *CEP192* | *EPHA5* | *KIAA0430* | *NLRP11* | *RAB40AL* | *STAC2* | *ZNF546* |
| *ALDH3B1* | *CFAP157* | *ESYT3* | *KIAA1217* | *NOP2* | *RAD54L2* | *SULT4A1* | *ZNF573* |
| *ALDH4A1* | *CFAP70* | *ETNPPL* | *KIAA1429* | *NOS2* | *RARG* | *SUN5* | *ZNF577* |
| *ANKHD1* | *CFL2* | *EXT2* | *KNTC1* | *NPAS2* | *RBM48* | *SYK* | *ZNF615* |
| *ANKHD1-EIF4EBP3* | *CHD1L* | *FAHD1* | *KPNA3* | *NPHP1* | *RBMS3* | *SYNRG* | *ZNF630-AS1* |
| *ANKLE2* | *CHD7* | *FAHD2B* | *L2HGDH* | *NSD2* | *RBPJ* | *SYT15* | *ZNF716* |
| *ANKRD6* | *CIT* | *FAM134B* | *LAMB1* | *NUTM1* | *RCC2* | *TAS1R3* | *ZNF746* |
| *ANO3* | *CLCA4* | *FAM227B* | *LAMB2* | *OIT3* | *RDH12* | *TAS2R41* | *ZNF781* |
| *ANO4* | *CMYA5* | *FAM98B* | *LARGE1* | *OR10W1* | *RGS22* | *TAS2R50* | *ZP2* |
| *ANPEP* | *CNKSR3* | *FANCL* | *LDOC1L* | *OR1S1* | *RHBDD1* | *TBC1D31* | *ZPBP2* |
| *APEH* | *COBL* | *FGD6* | *LHPP* | *OR5K1* | *RLN1* | *TBX15* | *ZZZ3* |

Sup Table 1: Genes with relevant genetic variants according the prioritization criteria.
